# The bounded coalescent model: conditioning a genealogy on a minimum root date

**DOI:** 10.1101/2022.01.04.474909

**Authors:** Jake Carson, Alice Ledda, Luca Ferretti, Matt Keeling, Xavier Didelot

**Affiliations:** Mathematics Institute, University of Warwick, United Kingdom; HCAI, Fungal, AMR, AMU & Sepsis Division, UK Health Security Agency, United Kingdom; Big Data Institute, University of Oxford, United Kingdom; Department of Statistics and School of Life Sciences, University of Warwick, United Kingdom

**Keywords:** coalescent model, most recent common ancestor, heterochronous sampling, phylogenetics, phylodynamics

## Abstract

The coalescent model represents how individuals sampled from a population may have originated from a last common ancestor. The bounded coalescent model is obtained by conditioning the coalescent model such that the last common ancestor must have existed after a certain date. This conditioned model arises in a variety of applications, such as speciation, horizontal gene transfer or transmission analysis, and yet the bounded coalescent model has not been previously analysed in detail. Here we describe a new algorithm to simulate from this model directly, without resorting to rejection sampling. We show that this direct simulation algorithm is more computationally efficient than the rejection sampling approach. We also show how to calculate the probability of the last common ancestor occurring after a given date, which is required to compute the probability of realisations under the bounded coalescent model. Our results are applicable in both the isochronous (when all samples have the same date) and heterochronous (where samples can have different dates) settings. We explore the effect of setting a bound on the date of the last common ancestor, and show that it affects a number of properties of the resulting phylogenies. All our methods are implemented in a new R package called BoundedCoalescent which is freely available online.

## 1 Introduction

The coalescent model is a stochastic process that describes the ancestry of a sample of individuals within a population (Kingman, 1982b,a). Conditioning the most recent common ancestor of the sample to be after a certain date results in a model called the “bounded coalescent” model and which was first mentioned in a model unifying gene duplication, loss and coalescence (Rasmussen and Kellis, 2012). In this context, the bound condition is used to deal with incomplete lineage sorting, which could cause a gene tree to be incongruent with the species tree (Maddison, 1997; Maddison and Knowles, 2006). Consequently, the bounded coalescent model is used in many multi-species coalescent models, to enforce the full coalescence of members of a same species before the speciation event (Mallo et al., 2016; Du et al., 2019; Hill et al., 2020; Li et al., 2021). The bounded coalescent model is also used in work on homologous recombination resulting in ancestral recombination graphs (Ferretti et al., 2013; Rasmussen et al., 2014). Furthermore, the bounded coalescent model is useful to perform pathogen transmission analysis from genetic data whilst accounting for within-host diversity (Didelot et al., 2014, 2017). In this case, setting a bound on the coalescent process equates to assuming a complete transmission bottleneck, so that all pathogen lineages within an infected individual need to coalesce before the host became infected. This results in a simpler relationship between the transmission tree of who-infected-whom and the genealogy of the pathogen sampled from the infected individuals. Despite this increasingly frequent use of the bounded coalescent model in several different biological research fields, the consequences of imposing a minimum on the date of the last common ancestor have not been formally investigated. In particular, under the standard coalescent model the waiting times between coalescent events are independent whereas this is no longer the case in the bounded coalescent model (Rasmussen and Kellis, 2012) since an increase in a waiting time needs to be compensated by a decrease in other waiting times to satisfy the bound condition.

In this paper, we start by defining the bounded coalescent model formally in both isochronous and heterochronous sampling settings. In the former all individuals are sampled at the same time, so that the genealogy is ultrametric (all leaves have the same distance to the root), whereas in the latter the individuals are sampled at different times, so that the genealogy is not ultrametric (Rambaut, 2000). We show how the probability of any genealogy can be computed under the bounded coalescent model with a given effective population size and bound date. This requires to first compute the probability of having the bound property occurring under a standard unbounded coalescent model, and we show how this can be computed efficiently. We also present a new algorithm for the simulation of genealogies under the bounded coalescent model given the sample number and dates, the effective population size and the bound date. Our algorithm can simulate directly from the bounded coalescent model, unlike previously described approaches which used rejection sampling on trees simulated from the standard unbounded coalescent model (Didelot et al., 2014; Mallo et al., 2016). These new algorithms to calculate the probability of genealogies and to simulate directly under the bounded coalescent model are both useful to perform inference under the model. Finally, we investigate a number of properties of the genealogies arising from the bounded coalescent model and how they differ from the unbounded coalescent model.

## 2 Definitions and notations

Coalescent models are typically derived from forward-in-time population models, such as the Wright-Fisher model (Wright, 1931; Fisher, 1930), the Moran model (Moran, 1958) and the Cannings exchangeable model (Cannings, 1974). The Wright-Fisher model assumes that a population evolves through non-overlapping generations, with each individual in each generation having a random uniformly distributed ancestor in the previous generation, independent of the others. Under a constant population size *N*, the probability that two individuals have the same ancestor in the previous generation is 1*/N*, so that the number of generations back-in-time until a common ancestor is found follows a Geometric distribution with mean *N*. This can be converted to real time by multiplying the number of generations by the generation interval *T*_*g*_, resulting in the effective population size *N*_*e*_ = *NT*_*g*_. Where *N*_*e*_ is sufficiently large, we can instead use the Kingman coalescent model (Kingman, 1982b,a), which replaces the Geometric distribution with an Exponential distribution and provides a continuous-time equivalent.

Under the standard coalescent model, the waiting time for a coalescence with *a* lineages is

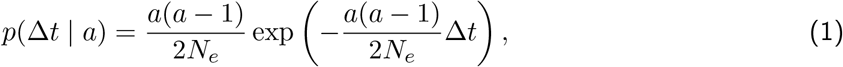

since each pair of lineages coalesces at rate 1*/N*_*e*_ and there are 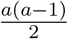 pairs of lineages.

In the most commonly used coalescent setting, *L* samples are taken simultaneously (isochronous) so that their ancestral process is simply made of the *L* − 1 coalescent events occurring back in time until the most recent common ancestor (MRCA) of the *L* samples is found. The probability of the coalescent dates in this isochronous setting can therefore be computed as the product of *L* − 1 terms given by Equation (1).

Here we consider a frequently used extension of this setting, in which the leaves are taken at different dates (heterochronous) *t*_1_ *< t*_2_ *<* … *< t*_*K*_ (Drummond et al., 2002, 2003). We define *L*_*k*_ as the number of leaves taken at date 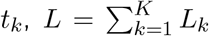 as the total number of leaves, and *τ*_1_ *< τ*_2_ *<* … *< τ*_*L*−1_ as the coalescent node times. Here and throughout this manuscript time is measured in the forward direction, so that for example *τ*_1_ is the date of the MRCA, and *t*_*K*_ is the date of the most recent sample. Furthermore we define the number of extant lineages at time *t* as

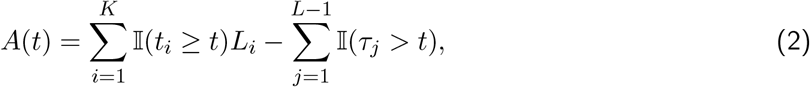

so that if *t* is a coalescence time, *A*(*t*) is the number of lineages that could have coalesced. These definitions are illustrated using an example phylogeny in Figure 1. Note that the isochronous case is a special case of the heterochronous case in which *K* = 1 and *L*_1_ = *L*.

**Figure 1:**
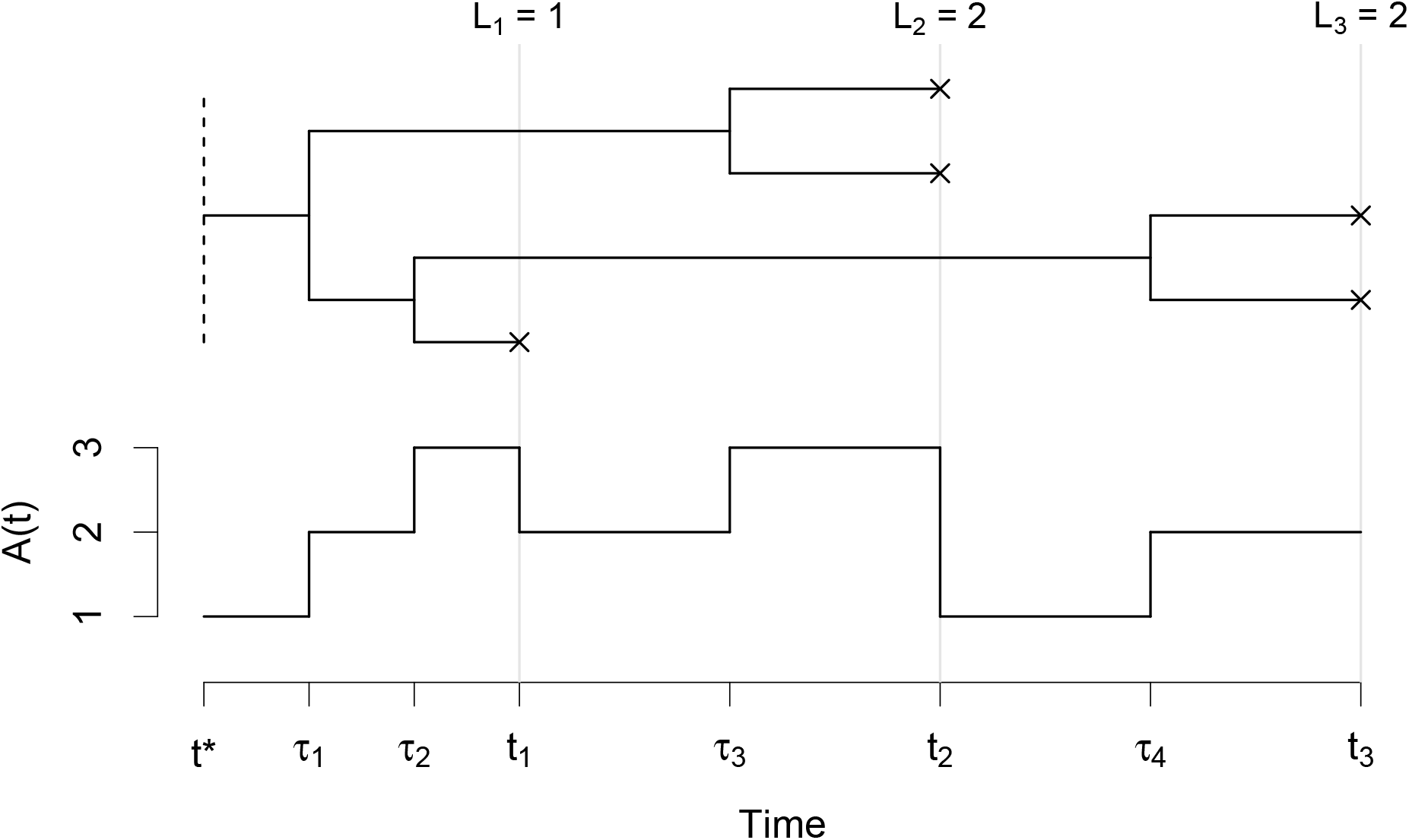
An example phylogeny in the heterochronous setting. Leaves are taken at times *t*_1_, *t*_2_, and *t*_3_, numbering *L*_1_ = 1, *L*_2_ = 2, and *L*_3_ = 2 leaves respectively (indicated by *×*). Lineages coalesce at times *τ*_1_, *τ*_2_, *τ*_3_, and *τ*_4_. *A*(*t*) is the number of lineages that may coalesce at time *t*. In the bounded coalescent model all lineages must coalesce by time *t*^*^ (dashed line), such that *A*(*t*^*^) = 1.

Letting *s*_1_, …, *s*_*K*+*L*−1_ be the ordered union of the leaves and coalescent node times, and *D*_*k*_ = (*t*_*k*_, *L*_*k*_) denote the combined sampling information, the probability of the ancestor dates in a heterochronous setting can be calculated as follows (Drummond et al., 2002):

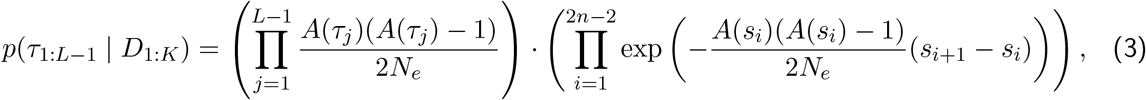

where we use the notations *τ*_1:*L*−1_ = *τ*_1_, …, *τ*_*L*−1_ and *D*_1:*K*_ = *D*_1_, …, *D*_*K*_.

In the bounded coalescent model we have the additional requirement that the lineages must coalesce to their MRCA before some specified bound time *t*^*^, so that *τ*_1_ *> t*^*^. We therefore write the probability of ancestor dates under the bounded coalescent model *p*(*τ*_1:*L*−1_ | *D*_1:*K*_, *τ*_1_ *> t*^*^) as opposed to *p*(*τ*_1:*L*−1_ | *D*_1:*K*_) for the unbounded coalescent model. When this condition is satisfied for two trees, their probability ratio under the bounded coalescent model should be the same as their probability ratio under the standard (unbounded) coalescent model:

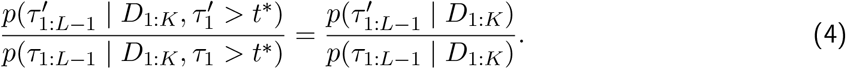

The probability of the ancestor dates under the bounded coalescent model can be rewritten using Bayes rule:

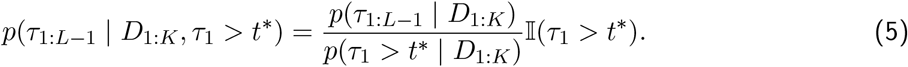

Note that the numerator is given by Equation (3). In other words the bounded coalescent model adds a normalising constant (denominator in Equation (5)) equal to the probability of all lineages coalescing, *p*(*τ*_1_ *> t*^*^ | *D*_1:*K*_), which we call the “bound probability” and which can be computed as shown in the next section. The Equation (5) clearly satisfies the ratio condition in Equation (4) since the denominators are the same and cancel out. Finally, we note that when *t*^*^ → −∞ the bound condition is always satisfied and the bound probability becomes one. In this case the bounded coalescent model reduces to the unbounded coalescent model, which in other words means that the bounded coalescent model is an extension of the unbounded coalescent model.

## 3 Bound probability

In an isochronous setting the bound probability is equal to the probability that *L* lineages coalesce into one within the time interval between sampling and *t*^*^. This probability is directly derived from the density function of the time to the MRCA in the standard coalescent model, which can be computed using matrix operations on the Kingman’s Markov chain (Tavaré, 1984), using a Laplace transform (Takahata and Nei, 1985) or using a convolution of exponential distributions (Wakeley, 2009) from the coalescent waiting times in Equation (1). Slightly more generally, the probability that *i* lineages coalesce down to *j* ≤ *i* lineages in time Δ*t* has also been computed before (Tavaré, 1984; Nordborg, 1998; Wakeley, 2009) and can be expressed as

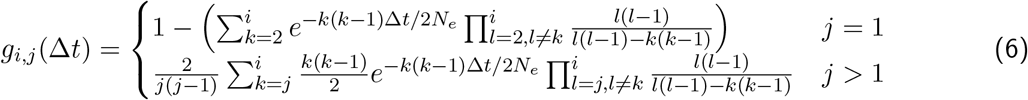

Hence the bound probability for the isochronous case is simply *p*(*τ*_1_ *> t*^*^ | *t*_1_, *L*) = *g*_*L*,1_(*t*_1_ − *t*^*^).

The heterochronous setting is more complex as lineages do not monotonically decrease with every coalescence. To simplify notation, let *A*_*k*_ = *A*(*t*_*k*_) denote the number of lineages at time *t*_*k*_. Note the following Markov property for the unbounded coalescent process

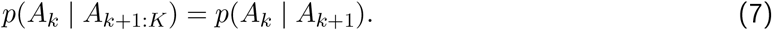

Consequently, we can express the number of lineages at a discrete set of time points as a hidden Markov model (HMM). We can then determine the bound probability using the forward algorithm (Rabiner, 1989; Zucchini and MacDonald, 2009).

The forward algorithm provides the set of probabilities *p*(*A*_*k*_ = *a*_*k*_), which are the probabilities of having *a*_*k*_ lineages at time *t*_*k*_ under the standard coalescent model. The algorithm is initialised at time *t*_*K*_ with *p*(*A*_*K*_ = *L*_*K*_) = 1, and iterates through *k* = *K* − 1, *K* − 2, …, 1 in order to evaluate

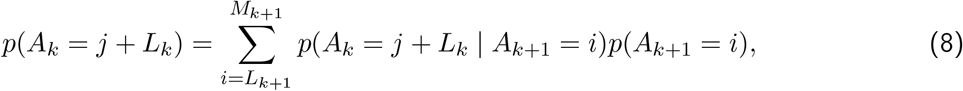

for *j* = 1, …, *M*_*k*+1_, where 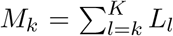 is the maximum number of lineages that can exist at time *t*_*k*_. The transition probabilities *p*(*A*_*k*_ = *j* + *L*_*k*_ | *A*_*k*+1_ = *i*) correspond to the probability in the standard coalescent model that *i* lineages coalesce down to *j* lineages in a time interval *t* = *t*_*k*+1_ − *t*_*k*_ as given in Equation (6), that is *p*(*A*_*k*_ = *j* + *L*_*k*_ | *A*_*k*+1_ = *i*) = *g*_*i,j*_(*t*_*k*+1_ − *t*_*k*_). Since *L*_*k*_ additional leaves are sampled at time *t*_*k*_, this results in *j* + *L*_*k*_ lineages.

The forward algorithm is terminated at the bound time *t*^*^. Defining *A*^*^ as the number of lineages at the bound time, calculation of the forward probabilities *p*(*A*^*^) follow Equation (8) but without new leaves being added:

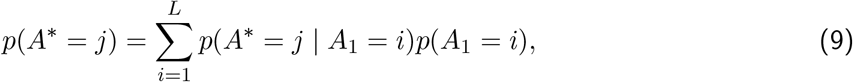

for *j* = 1, …, *L*, where *p*(*A*^*^ = *j* | *A*_1_ = *i*) = *g*_*i,j*_(*t*_1_ − *t*^*^). The bound probability is then given by *p*(*A*^*^ = 1).

This concludes the calculation of the bound probability in the heterochronous case. This quantity is of interest by itself, but also and perhaps more importantly it allows the calculation of the probability of sampling a tree under the bounded coalescent model by applying Equation (5). This calculation allows inference to be performed under the bounded coalescent model, either using maximum-likelihood or in a Bayesian framework.

## 4 Direct sampling

A straightforward approach to simulate realisations of the bounded coalescent model is to use rejection sampling (Didelot et al., 2014; Mallo et al., 2016). This involves simulating from the standard coalescent model and keeping only those simulations in which *τ*_1_ *> t*^*^. This rejection sampling approach can also be used to estimate the bound probability, since acceptance happens with probability equal to the bound probability. However, rejection sampling will be inefficient especially if lineages are sampled close to the bound relative to the effective population size, i.e. if 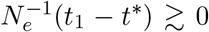. In this case the bound probability is small and therefore so is the acceptance probability of the rejection sampler. Here we introduce a direct sampler for the bounded coalescent model that does not suffer this limitation.

The direct sampling approach proceeds through the following five steps:

Step 1 Use the forward filtering backward sampling (FFBS) algorithm to sample the number of lineages *A*_1:*K*_ at times *t*_1:*K*_ conditioned on the bound.

Step 2 From the sampled *A*_1:*K*_ derive the number of coalescence events in the time intervals (*t*^*^, *t*_1_) and (*t*_*k*_, *t*_*k*+1_) for *k* = 1, …, *K* − 1.

Step 3 For intervals containing multiple coalescence events, subdivide the time interval and sample the number of coalescence events for each subinterval. Repeat this step until all *L* − 1 coalescence events are constrained by unique, non-overlapping time intervals.

Step 4 Use inverse transform sampling to sample the (constrained) coalescence times.

Step 5 Working backwards in time, sample a pair of lineages for each coalescence event.

Further details of each step are discussed below. We also describe how to calculate the probability *p*(*τ*_1:*L*−1_ | *D*_1:*K*_, *τ*_1_ *> t*^*^) concurrently with the sampling approach, allowing the sampler to be efficiently utilised within an inferential framework. For example it allows the use of the sampler as a proposal distribution in a Markov Chain Monte-Carlo or Importance Sampling algorithm, since in both cases the probability of the proposed tree would be needed.

### 4.1 Step 1

In Section 3 we describe how the number of lineages *A*^*^, *A*_1:*K*_ can be expressed as a HMM. By treating *A*^*^ = 1 as an ‘observation’, we can simulate values of *A*_1:*K*_ conditioned on *A*^*^ = 1 using the FFBS algorithm. FFBS uses a two-step recursion: a forward recursion to calculate the forward probabilities, and a backward recursion to generate samples. The forward recursion is the forward algorithm described in Section 3, so here we focus on the backward recursion for sampling.

The backward recursion is initiated by calculating the probabilities

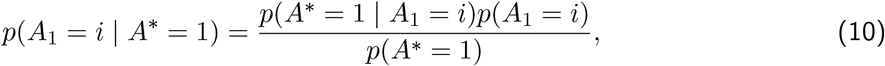

for *i* = 1, …, *L*. The terms *p*(*A*_1_ = *i*) and *p*(*A*^*^ = 1) are probabilities calculated in the forward algorithm, and the transition density is given by Equation (6), i.e. *p*(*A*^*^ = 1 | *A*_1_ = *i*) = *g*_*i*,1_(*t*_1_ − *t*^*^). Once these probabilities have been calculated, the number of lineages *a*_1_ is sampled according to the probabilities *p*(*A*_1_ = *a*_1_ | *A*^*^ = 1). The backward recursion then iterates through *k* = 2,.., *K*, calculating the probabilities

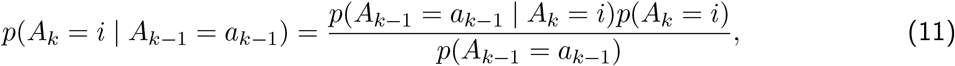

and sampling a value *a*_*k*_ accordingly. Note again that the transition density must account for the addition of leaves, i.e. *p*(*A*_*k*−1_ = *j* + *L*_*k*−1_ | *A*_*k*_ = *i*) = *g*_*i,j*_(*t*_*k*_ − *t*_*k*−1_).

The calculation of *p*(*τ*_1:*L*−1_ | *D*_1:*K*_, *τ*_1_ *> t*^*^) is initialised by setting

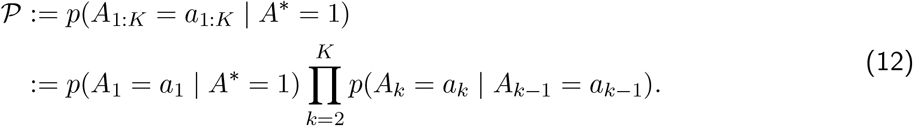

We use 𝒫 to emphasise that this is a partial calculation, and will be updated in the following steps.

### 4.2 Step 2

Having sampled the number of lineages *a*_1:*K*_ we derive the number of coalescence events between successive time points. Define *c*_*,1_ as the number of coalescence events in the interval (*t*^*^, *t*_1_), and *c*_*k*−1,*k*_ as the number of coalescence events in the interval (*t*_*k*−1_, *t*_*k*_). Then

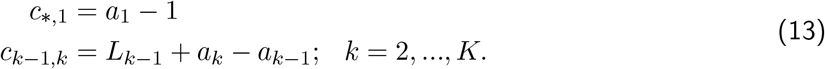

### 4.3 Step 3

In order to sample coalescence times in Step 4, we require that each coalescence event is constrained within a unique, non-overlapping time interval. Hence, for any *c*_*k*−1,*k*_ ≥ 2, we need to partition the corresponding time interval until the coalescence events are separated. Here, we bisect the interval (*t*_*k*−1_, *t*_*k*_) and sample the number of coalescent events in the subintervals (*t*_*k*−1_, *t*_*k*−0.5_) and (*t*_*k*−0.5_, *t*_*k*_), where *t*_*k*−0.5_ = 0.5(*t*_*k*−1_ + *t*_*k*_). This is achieved by sampling the number of lineages *a*_*k*−0.5_ at the newly added time point *t*_*k*−0.5_ according to

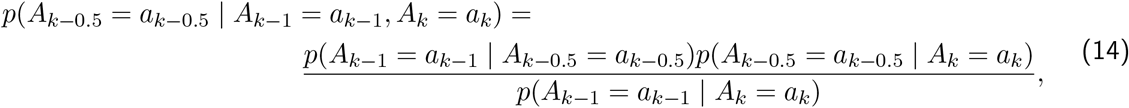

and then deriving the number of coalescence events *c*_*k*−1,*k*−0.5_ and *c*_*k*−0.5,*k*_ as in Step 2. The probability of sampling *a*_*k*−0.5_ is incorporated into the calculation of *p*(*τ*_1:*L*−1_ | *D*_1:*K*_, *τ*_1_ *> t*^*^):

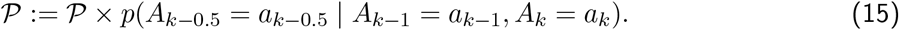

This process is also applied to the interval (*t*^*^, *t*_1_) if *c*_*,1_ ≥ 2, and any newly generated subintervals until we have at most one coalescence event in any defined time interval.

### 4.4 Step 4

Since each coalescence event is constrained by a unique non-overlapping time interval, we can sample the coalescence times using inverse transform sampling. From the previous steps, assume that the coalescence event occurs in the interval (*t*_*k*−1_, *t*_*k*_), with *a*_*k*_ lineages at time *t*_*k*_ and *a*_*k*−1_ = *a*_*k*_ *–* 1 lineages at time *t*_*k*−1_. The density function the coalescence time *τ* within the interval is

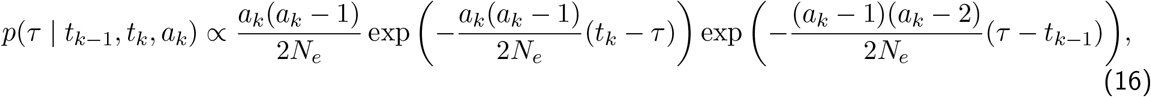

where the first term corresponds to the density of coalescence time under the standard coalescent model, and the second term corresponds to the probability that no further coalescence events occur in the interval (*t*_*k*−1_, *τ*). Collecting the *τ* terms gives

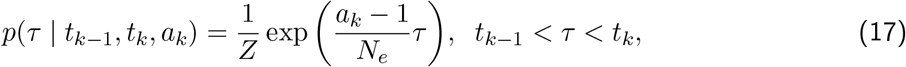

where *Z* is the normalising constant, and is equal to

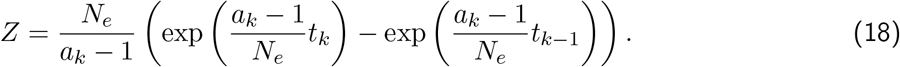

If *u* is a draw from a standard uniform distribution, we can compute *τ* using inverse transform sampling by solving

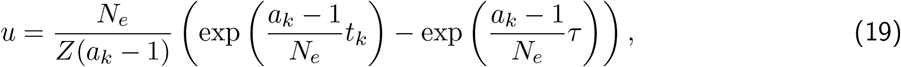

where the right side is the cumulative density function of *τ*. This gives

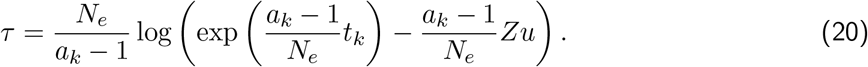

Each time a coalescence is sampled, the calculation of *p*(*τ*_1:*L*−1_ | *D*_1:*K*_, *τ*_1_ *> t*^*^) is updated using

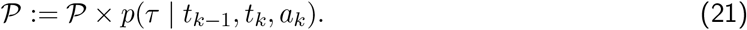

By using inverse transform sampling for each coalescence event, we obtain the full set of coalescence times *τ*_1:*L*−1_. The probability *P* at the end of this step is equal to that given by Equation (5), i.e. 𝒫 = *p*(*τ*_1:*L*−1_ | *D*_1:*K*_, *τ*_1_ *> t*^*^).

### 4.5 Step 5

The final step is to sample the topology of the tree. Here we simply iterate backwards in time through the coalescence events and sample two of the extant lineages, noting the ancestors for each coalescent node. The probability of a topology *G* conditional on the sampled *τ*_1:*L*−1_ is given by

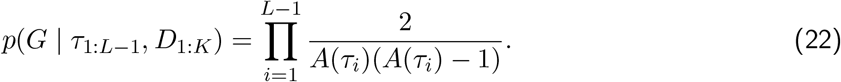

The probability density of the sampled tree under the bounded coalescent model is then given by

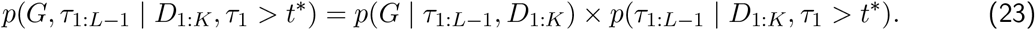

### 4.6 Validation and comparison with rejection sampling

#### 4.6.1 Bound probability

Here we consider an example, where leaves are added at times *t*_1_ = 0.0, *t*_2_ = 0.5, *t*_3_ = 1.0, *t*_4_ = 1.5, *t*_5_ = 2.0 and must coalesce by time *t*^*^ = −0.5. We use an effective population size of *N*_*e*_ = 1 so that pairs of lineages coalesce at rate 1*/N*_*e*_ = 1. We first used a rejection approach to simulate under these conditions. The rejection sampler required 427 371 simulations in order to obtain 10^6^ acceptances, which means that the bound probability is estimated to be 0.234.

Next we run the forward filter described in Section 3, giving the probabilities shown in Table 1. In particular we note that the bound probability of having a single lineage at the bound time *t*^*^ is 0.233, which is consistent with the bound probability estimated by the rejection sampler.

**Table 1:**
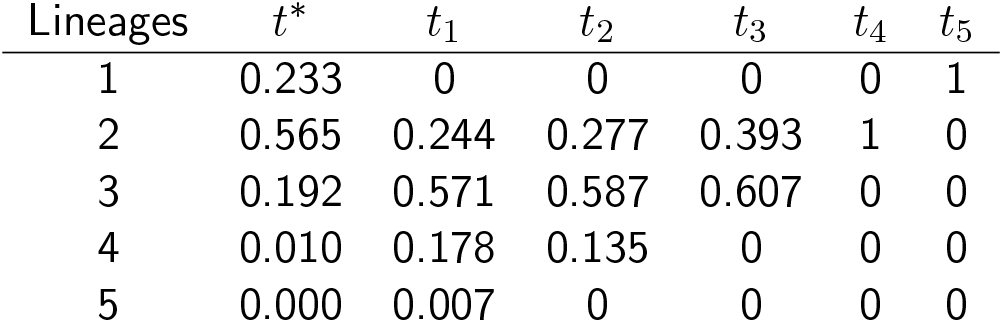
Forward filter probabilities in the numerical example (probabilities may not add to 1 due to rounding).

| Lineages | $t^*$ | $t_1$ | $t_2$ | $t_3$ | $t_4$ | $t_5$ |
| --- | --- | --- | --- | --- | --- | --- |
| 1 | 0.233 | 0 | 0 | 0 | 0 | 1 |
| 2 | 0.565 | 0.244 | 0.277 | 0.393 | 1 | 0 |
| 3 | 0.192 | 0.571 | 0.587 | 0.607 | 0 | 0 |
| 4 | 0.010 | 0.178 | 0.135 | 0 | 0 | 0 |
| 5 | 0.000 | 0.007 | 0 | 0 | 0 | 0 |

#### 4.6.2 A single run of the direct sampler

Step 1 of the direct sampler consists in running the forward algorithm to obtain the probabilities shown in Table 1, followed by backwards sampling conditioned on the bound, that is *A*^*^ = 1. We update the filtered probabilities for *A*_1_ using Equation (11), giving the probability vector (0.000, 0.411, 0.494, 0.093, 0.002)^T^, from which we sample. Say we sample *A*_1_ = 4, we iterate to time *t*_2_ and obtain the probabilities *p*(*A*_2_ = 3 | *A*_1_ = 4) = 0.736 and *p*(*A*_2_ = 4 | *A*_1_ = 4) = 0.264. Note that *p*(*A*_2_ = 2 | *A*_1_ = 4) = 0 since 2 leaves at time *t*_2_ can not result in 4 leaves at time *t*_1_. This sampling procedure continues until we have samples *a*_1_, …, *a*_5_.

In Step 2 we determine the number of coalescence events between time points. If in Step 1 we sample *a*_1_ = 4, *a*_2_ = 3, *a*_3_ = 3, *a*_4_ = 2, *a*_5_ = 1 we can determine that *c*_*,1_ = 3 coalescence events occur in the interval (*t*^*^, *t*_1_), and *c*_2,3_ = 1 in (*t*_2_, *t*_3_).

In Step 3 we further partition the time axis in order to separate each coalescence event. Here we would add a new time point at *t*_0.5_ = −0.25, and sample the extant number of lineages conditional on having a single lineage at time *t*^*^ and four lineages at time *t*_1_. Hence we may sample 0, 1, 2, or 3 coalescence events in the interval (*t*^*^, *t*_0.5_), with the remainder occurring in the interval (*t*_0.5_, *t*_1_). This partitioning continues until each coalescence events is constrained by a unique non-overlapping time interval, i.e. (−0.5, −0.25), (−0.25, −0.125), (−0.125, 0.0), (0.5, 1.0).

Step 4 is simply a matter of sampling the coalescence times, conditioned on the corresponding intervals.

Finally, in Step 5 we sample the lineages for each coalescence event. Starting with the latest event, we determine which lineages currently exist. For the interval (0.5, 1.0) these would be the lineages corresponding to leaves 3, 4, and 5. In the interval (−0.125, 0.0), these would be the lineages corresponding to leaves 1 and 2, as well as the resulting lineages from the previous coalescence event.

#### 4.6.3 Comparison of simulated trees using direct and rejection sampling

We obtain 10^6^ simulations from both the rejection sampling algorithm and the direct sampling algorithm as described above. In Figure 2 we compare the simulated coalescence times, and observe strong agreement between the two methods.

**Figure 2:**
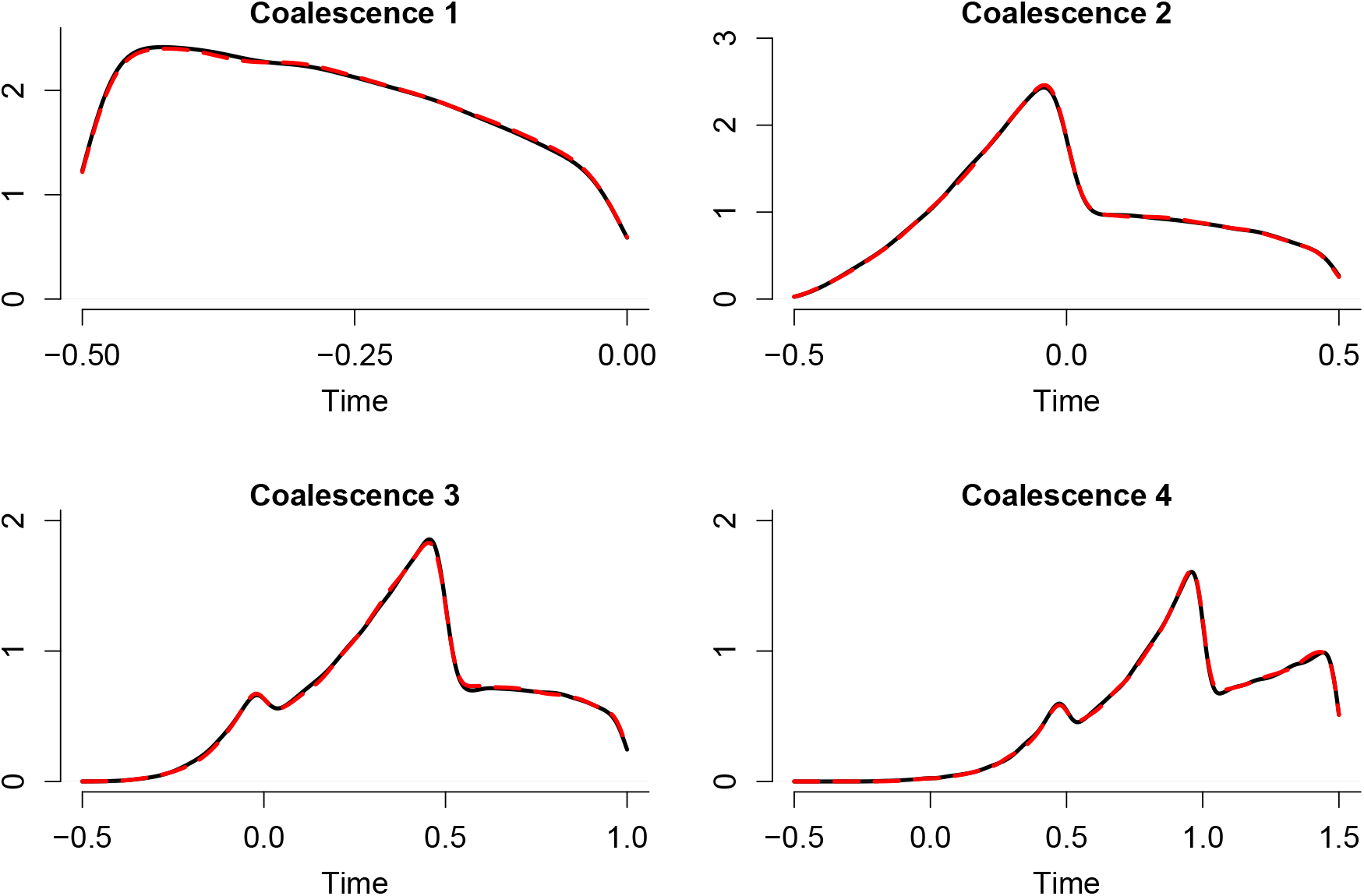
Estimates of the four coalescence times. The solid black line is the density obtained using direct sampling, and the dashed red line is the density obtained using rejection sampling.

We also compare the run times of the two algorithms as we vary the number of leaves. Keeping *t*^*^ = −0.5 and *N*_*e*_ = 1 fixed, we sample *L* leaves uniformly over the interval (0, 2) and obtain 10^6^ simulations from each algorithm. The run times for *L* = 2, …, 50 are shown in Figure 3. For a small number of leaves, both algorithms exhibit similar performance. However, as the number of leaves increases, the computational cost of rejection sampling increases much more rapidly than the direct sampling approach. This results from the smaller bound probabilities lowering the acceptance rate of the rejection sampler.

**Figure 3:**
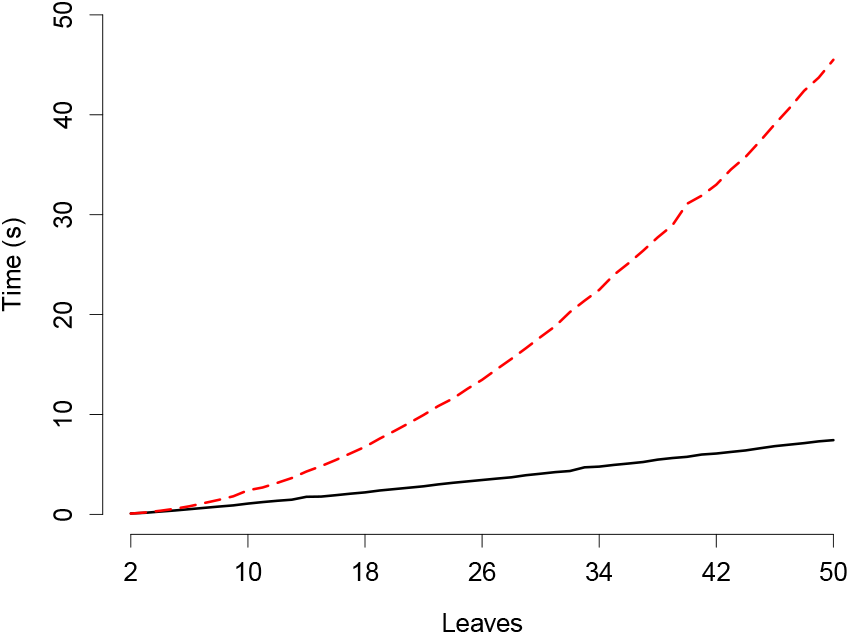
Comparison of run times for direct sampling and rejection sampling with varying number of leaves. The solid black line is the run time for direct sampling, and the dashed red line is the run time for rejection sampling. 10^6^ simulations are obtained in each instance.

## 5 Properties

In this section we investigate the effect of the bound on various properties of the tree, which can differ markedly from the standard coalescent model and alter the outcome of many standard analyses.

### 5.1 Dependencies between coalescent events

The bound induces dependencies between the waiting times of the coalescence events. As a demonstration, we consider the isochronous setting with *L* = 3, *t*_1_ = 0, and estimate the correlation between the two waiting times for bound times −5 ≤ *t*^*^ ≤ −0.01. This estimation is performed numerically by simulating 100 000 trees for each bound time. The results are shown in Figure 4. As the bound time increases, the correlation between waiting times becomes stronger. Since all lineages must coalesce by the bound time, if one waiting time is large then the other must be small. As the bound time decreases and the bounded coalescent model more strongly approximates the standard coalescent model, the correlation tends towards zero.

**Figure 4:**
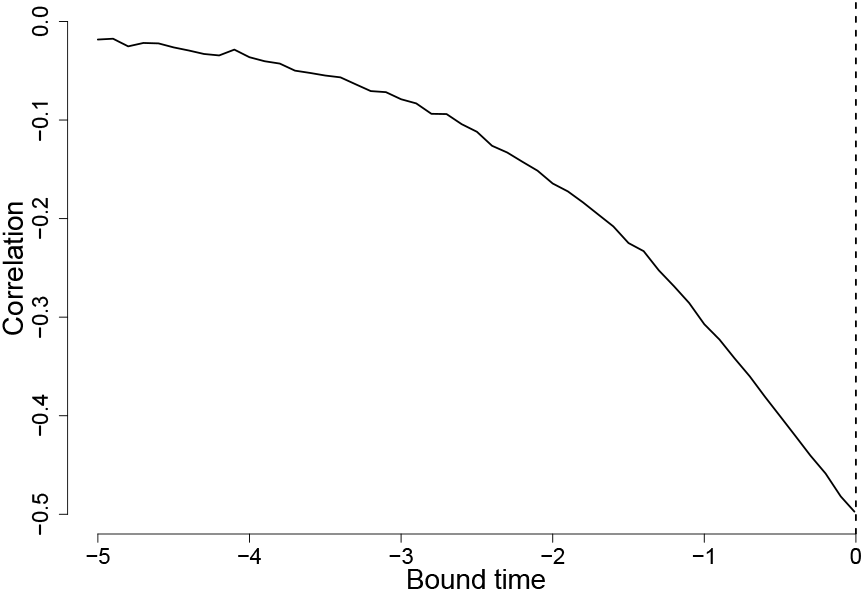
Correlation between the two waiting times when *L* = 3, *t*_1_ = 0 (dashed line) and *N*_*e*_ = 1.

Due to these dependencies, it is necessary to consider all leaves and lineages jointly when performing simulations or deriving probabilities under the bounded coalescent model. For instance, simulation under the standard coalescent model can be undertaken by adding leaves one by one to the growing tree, but this is not the case under the bounded coalescent model. To illustrate, we again consider the isochronous setting and estimate by simulation the average pairwise distance between leaves for *L* = 2, …, 50, while keeping *t*^*^ fixed at −0.5 (Figure 5a) and −2 (Figure 5b). In both cases, increasing the total number of leaves increases their average pairwise distance. Consequently, simulating a tree under the bounded coalescent model with *L* = 2 and adding a leaf is not equivalent to simulating a tree under the bounded coalescent model with *L* = 3.

**Figure 5:**
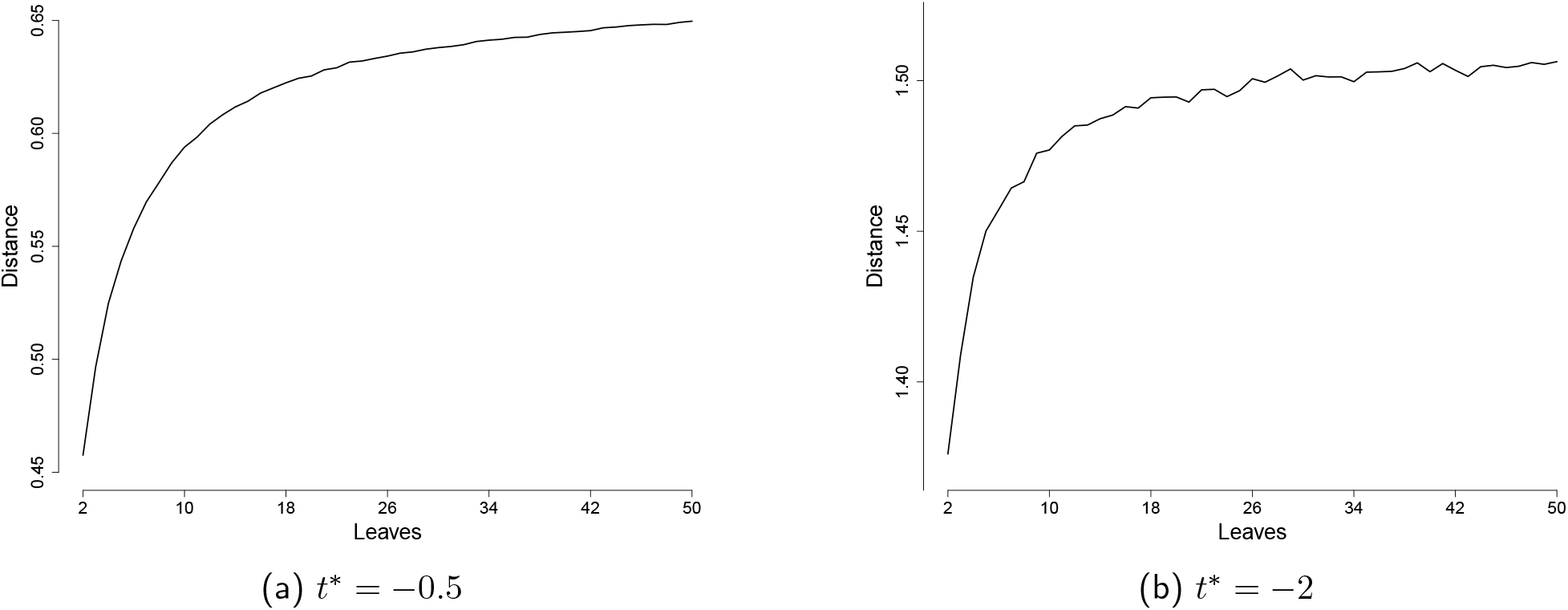
Average distance between two leaves for different values of *L* keeping *t*_1_ = 0 and *N*_*e*_ = 1 fixed.

### 5.2 Bound probability

For further exploration we consider the heterochronous setting where *L* leaves are sampled at evenly spaced times over the sampling interval [0, *t*_*L*_]. The isochronous setting can be recovered by setting *t*_*L*_ = 0. We investigate how properties of the simulated trees change as *L, t*_*L*_, and *t*^*^ are varied.

Estimates of the bound probability for varying *L, t*_*L*_, and *t*^*^ are shown in Figure 6. The value of the bound time *t*^*^ has the largest impact. As the bound time moves further into the past the bound probability tends towards one, even for large numbers of leaves and/or short sampling intervals. Sampled trees will strongly resemble those obtained under the standard coalescent model. On the other hand, as the bound time becomes more recent, the bound probability tends towards zero. In this region sampled trees will have very different properties than the standard coalescent model. For modest bound times, the choice of both *L* and *t*_*L*_ significantly impact the bound probability, which decreases for increasing *L* and decreasing *t*_*L*_. The largest changes are observed for small *L* and *t*_*L*_, and asymptotic behaviour is observed for large *L* and *t*_*L*_.

**Figure 6:**
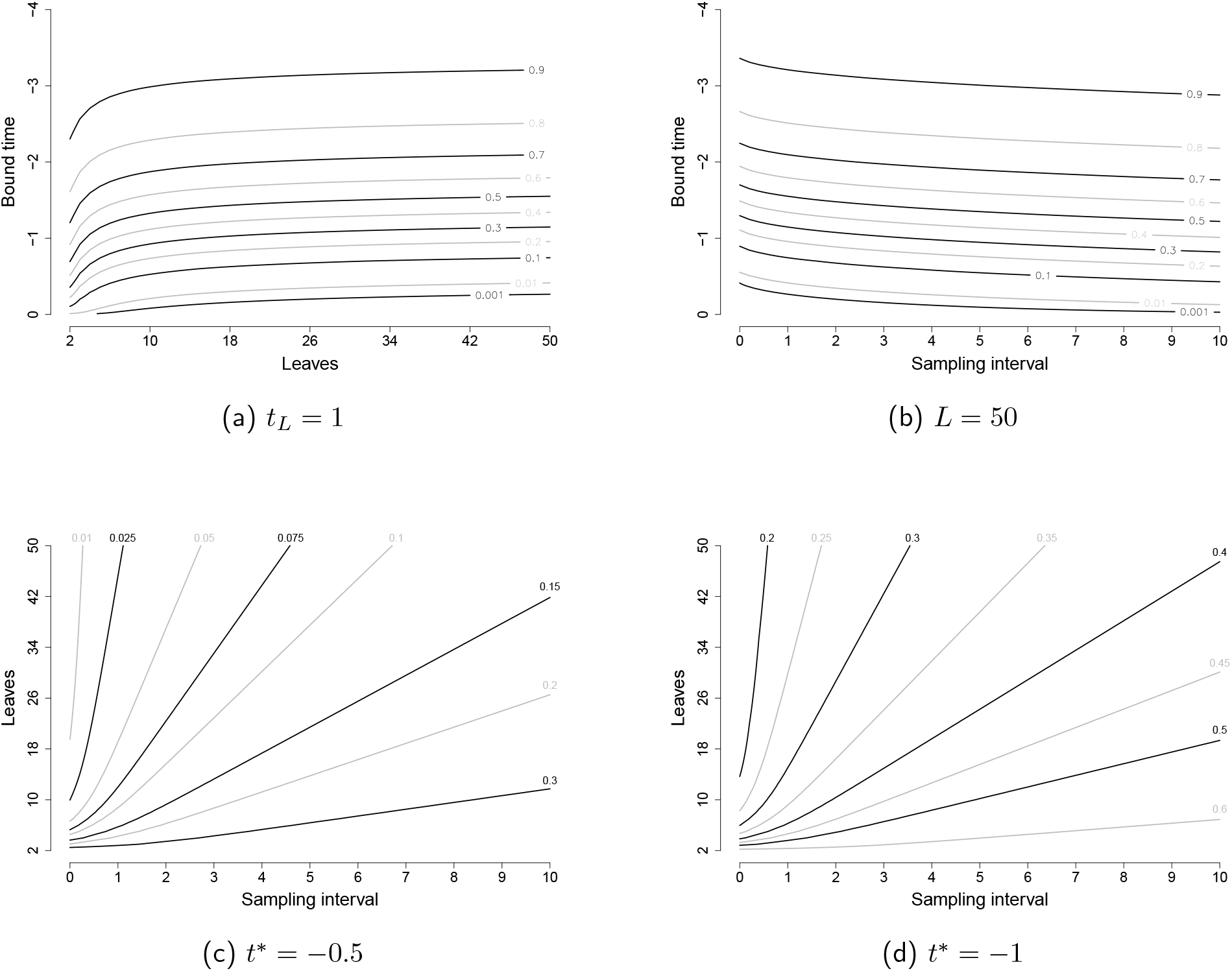
Bound probabilities as the bound time and number of leaves vary in the heterochronous example. (a) uses a sampling interval of 1, (b) uses 50 leaves, (c) uses a bound time of −0.5, and (d) uses a bound time of −1.

### 5.3 Tree summary statistics

Figure 7 shows how several summary statistics of bounded coalescent trees change as the bound time is altered. In particular we consider the distributions of the time of the MRCA (TMRCA), the total branch length, the average pairwise distance between the leaves, and the ratio between the average terminal branch length to the average internal branch length (starlikeness). The total number of leaves is fixed at *L* = 50 in Figure 7, and we consider three sampling intervals, *t*_*L*_ = 0 (isochronous), *t*_*L*_ = 1, and *t*_*L*_ = 10. Within these three configurations, we consider the four bound times *t*^*^ = −∞, *t*^*^ = −2, *t*^*^ = −1, and *t*^*^ = −0.5, and sample 1000 trees in each of the twelve cases to approximate the resulting distributions.

**Figure 7:**
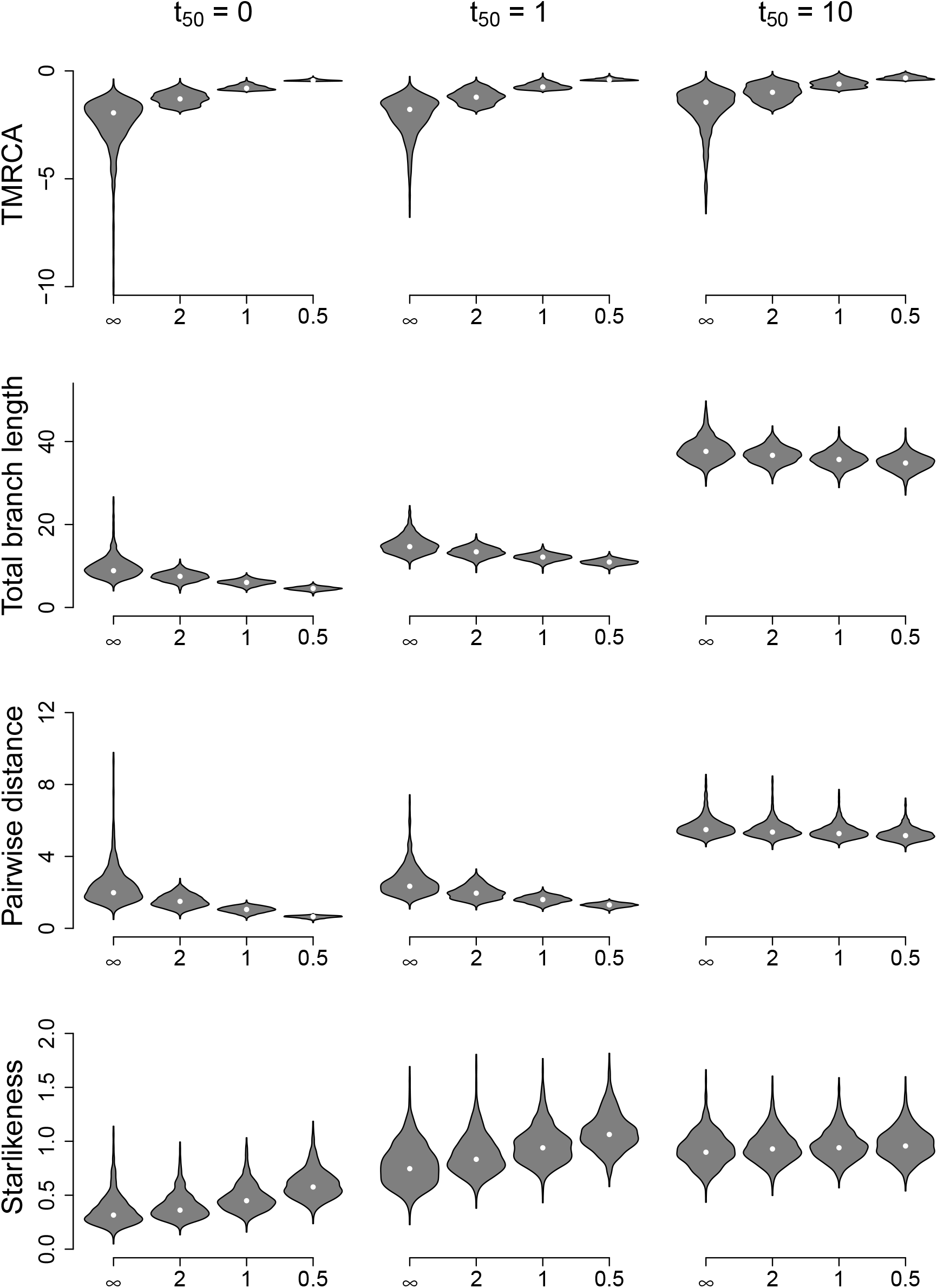
Properties of the bounded coalescent model with 50 leaves and sampling intervals of length 0 (isochronous), 1, and 10. Mean values are shown as white circles.

Since all lineages are required to coalesce before the bound time, providing a lower bound for TMRCA, the bounded coalescent model leads to more recent values of TMRCA. As the bound time increases, values of TMRCA become increasingly concentrated near the bound time. Increasing the bound time tends to reduce the total branch length. This results from requiring all lineages to coalesce by the bound time, which gives an upper limit of ((*t*_*L*_/2) − *t*^*^)*L*.

As with the total branch length, the bounded coalescent model places an upper bound on the average pairwise distance, which is (*L* + 1)/(3(*L* − 1)). This causes the average pairwise distance to decrease as the bound time increases. Finally, the starlikeness increases as the bound time increases.

### 5.4 Phylodynamics

The starlikeness of a tree is often an indication of past population size growth (Slatkin and Hudson, 1991; den Bakker et al., 2008; Volz et al., 2009). Therefore, the fact that this tree summary statistic depends on the bound time (Figure 7) suggests that the presence of a bound could confound phylodynamic inference studies, which are aimed at reconstruct past population size dynamics given genetic or phylogenetic data (Nee et al., 1995; Pybus et al., 2000; Ho and Shapiro, 2011).

To illustrate this, Figure 8 shows two examples of skyline plots in the isochronous setting and two examples in the heterochronous setting. All skyline plots were computed using Bayesian nonparametric phylodynamic reconstruction (Palacios and Minin, 2012) as implemented in the R package phylodyn (Karcher et al., 2017). In all four examples the population size was incorrectly inferred to have grown significantly. In examples (a) and (c) this was caused by a high effective population size relative to the bound time, so that the bound conditions forces coalescence to happen before it would normally do, whereas in examples (b) and (d) the same occurred due to relatively recent bound times. These results do not invalidate the principles of phylodynamic inference, which assume that there is no bound on the root date, but warn against its application in situations where a bound is present.

**Figure 8:**
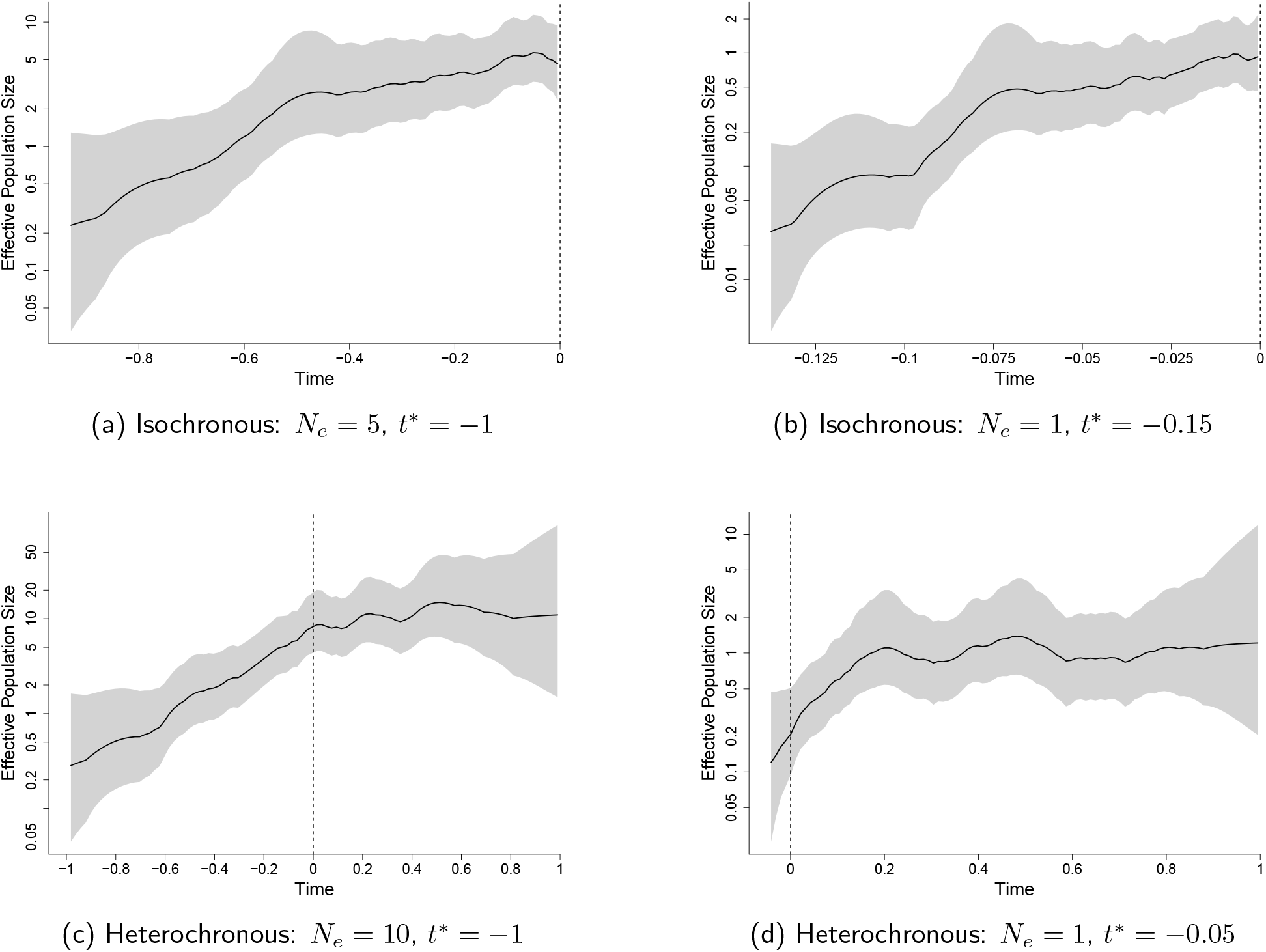
Skyline plots with *L* = 50. The solid lines show the median population estimates, and the grey regions show 95% credible intervals. The plotted range indicates the bound time and the latest leaf. The dashed line shows the earliest sampled leaf.

## 6 Implementation

We implemented the algorithms and methods described in this paper into a new R package called *BoundedCoalescent* which is available at https://github.com/DrJCarson/BoundedCoalescent. This package includes functions to calculate the bound probability, to calculate the probability density of a tree under the bounded coalescent model, and to simulate trees under the bounded coalescent model. Most of the code was written in C++ and integrated into the R package using Rcpp (Eddelbuettel and François, 2011; Eddelbuettel, 2013). The R package ape was used to store, manipulate and visualise phylogenetic trees (Paradis and Schliep, 2019).

## 7 Discussion

In this paper we have presented a formal description of the bounded coalescent model (Rasmussen and Kellis, 2012), an extension of the standard coalescent model in which all lineages are constrained to find a common ancestor by a predefined date. We have shown how to calculate the probability of the bound constraint happening by chance, which is useful to calculate the probability of a given phylogeny under the bounded coalescent model with a given bound date. We have also described a method to directly sample phylogenies under the bounded coalescent model, and used this to explore the properties of the model and the effect of the conditioning.

The standard coalescent framework can be extended in many ways (Donnelly and Tavare, 1995; Fu and Li, 1999; Rosenberg and Nordborg, 2002), for example to allow for variations in the population size (Griffiths and Tavare, 1994), geographical structure (Notohara, 1990), recombination (Hudson, 1990) and selection (Krone and Neuhauser, 1997). All of these extensions are in principle compatible with the conditioning imposed by the bounded coalescent model. For example, an ancestral recombination graph could be simulated, for which efficient methods have been developed (McVean and Cardin, 2005), and rejection sampling could be applied to ensure that the bound condition is met. However, it is unclear under which of these coalescent framework extensions a direct sampling method can be devised or an efficient method to calculate the probability of realisations.

The bounded coalescent model is of special interest for applying coalescent theory in infectious disease epidemiology. This includes the need to have full coalescence of lineages within a host before that host become infected, if we assume a complete transmission bottleneck (Didelot et al., 2014, 2017). The effect of a complete transmission bottleneck becomes more important as data on within-host diversity are increasingly being used to infer who infected whom (De Maio et al., 2018; Wymant et al., 2018). An alternative approach to the bounded coalescent model in order to enforce a complete transmission bottleneck is to start the within-host population with an effective population size of zero, with subsequent growth, so that the coalescence rate is close to infinity just after infection. For example, a within-host linear growth model starting at zero was used as part of a method for simultaneous inference of phylogenetic and transmission trees (Klinkenberg et al., 2017). Similarly, a recently proposed model on clonal expansion has each expansion starting with a size of zero to ensure they initially correspond to a single lineage (Helekal et al., 2021). Finally, the coalescent process is sometimes equated with the transmission process by assuming that the within-host population size is negligible, so that coalescent times correspond to transmission events (Volz et al., 2009; Frost and Volz, 2010). In this framework, considering that incidence is proportional to prevalence, as is the case in many infectious disease epidemiology models, leads to an effective population size proportion to the number of infected individuals minus one (Volz, 2012; Volz and Didelot, 2018). Consequently, the bound condition is enforced by having an effective population size of zero at the time when an outbreak is seeded with a single index case.

## Acknowledgements

We acknowledge funding from the National Institute for Health Research (NIHR) Health Protection Research Unit in Genomics and Enabling Data (grant number NIHR200892).

## References

den Bakker, H.C., Didelot, X., Fortes, E.D., Nightingale, K.K., Wiedmann, M., 2008. Lineage specific recombination rates and microevolution in Listeria monocytogenes. BMC Evol. Biol. 8, 277. doi:10.1186/1471-2148-8-277.

Cannings, C., 1974. The latent roots of certain Markov chains arising in genetics: a new approach, I. Haploid models. Adv. Appl. Probab. 6, 260–290. doi:10.2307/1426293.

De Maio, N., Worby, C.J., Wilson, D.J., Stoesser, N., 2018. Bayesian reconstruction of transmission within outbreaks using genomic variants. PLOS Comput. Biol. 14, e1006117. doi:10.1371/journal.pcbi.1006117.

Didelot, X., Fraser, C., Gardy, J., Colijn, C., 2017. Genomic infectious disease epidemiology in partially sampled and ongoing outbreaks. Mol. Biol. Evol. 34, 997–1007. doi:10.1093/molbev/msw275.

Didelot, X., Gardy, J., Colijn, C., 2014. Bayesian inference of infectious disease transmission from whole-genome sequence data. Mol. Biol. Evol. 31, 1869–1879. doi:10.1093/molbev/msu121.

Donnelly, P., Tavare, S., 1995. Coalescents and genealogical structure under neutrality. Annu. Rev. Genet. 29, 401–21. doi:10.1146/annurev.ge.29.120195.002153.

Drummond, A.J., Nicholls, G.K., Rodrigo, A.G., Solomon, W., 2002. Estimating mutation parameters, population history and genealogy simultaneously from temporally spaced sequence data. Genetics 161, 1307–1320. doi:10.1093/genetics/161.3.1307.

Drummond, A.J., Pybus, O.G., Rambaut, A., Forsberg, R., Rodrigo, A.G., 2003. Measurably evolving populations. Trends Ecol. Evol. 18, 481–488. doi:10.1016/S0169-5347(03)00216-7.

Du, P., Ogilvie, H.A., Nakhleh, L., 2019. Unifying gene duplication, loss, and coalescence on phylogenetic networks, in: Lect. Notes Comput. Sci.. Springer International Publishing. volume 11490 LNBI, pp. 40–51. doi:10.1007/978-3-030-20242-2\_4.

Eddelbuettel, D., 2013. Seamless R and C++ integration with Rcpp. Springer, New York. doi:10.1007/978-1-4614-6868-4.

Eddelbuettel, D., François, R., 2011. Rcpp: seamless R and C++ integration. J. Stat. Softw 40, 1–18. doi:10.18637/jss.v040.i08.

Ferretti, L., Disanto, F., Wiehe, T., 2013. The effect of single recombination events on coalescent tree height and shape. PLoS One 8, e60123. doi:10.1371/journal.pone.0060123.

Fisher, R.A., 1930. The genetical theory of natural selection. Clarendon Press. doi:10.5962/bhl.title.27468.

Frost, S.D.W., Volz, E.M., 2010. Viral phylodynamics and the search for an ‘effective number of infections’. Philos. Trans. R. Soc. B 365, 1879–1890. doi:10.1098/rstb.2010.0060.

Fu, Y.X., Li, W.H., 1999. Coalescing into the 21st century: An overview and prospects of coalescent theory. Theor. Popul. Biol. 56, 1–10. doi:10.1006/tpbi.1999.1421.

Griffiths, R.C., Tavare, S., 1994. Sampling theory for neutral alleles in a varying environment. Philos. Trans. R. Soc. B 344, 403–410. doi:10.1098/rstb.1994.0079.

Helekal, D., Ledda, A., Volz, E., Wyllie, D., Didelot, X., 2021. Bayesian inference of clonal expansions in a dated phylogeny. Syst. Bio. TBD, syab095. doi:10.1093/sysbio/syab095.

Hill, M., Legried, B., Roch, S., 2020. Species tree estimation under joint modeling of coalescence and duplication: sample complexity of quartet methods. arXiv, 2007.06697.

Ho, S.Y.W., Shapiro, B., 2011. Skyline-plot methods for estimating demographic history from nucleotide sequences. Mol. Ecol. Resour. 11, 423–434. doi:10.1111/j.1755-0998.2011.02988.x.

Hudson, R.R., 1990. Gene genealogies and the coalescent process. Oxford Surv. Evol. Biol. 7, 1–44.

Karcher, M.D., Palacios, J.A., Lan, S., Minin, V.N., 2017. PHYLODYN: an R package for phylodynamic simulation and inference. Mol. Ecol. Resour. 17, 96–100. doi:10.1111/1755-0998.12630.

Kingman, J.F.C., 1982a. On the genealogy of large populations. J. Appl. Probab. 19, 27–43. doi:10.2307/3213548.

Kingman, J.F.C., 1982b. The coalescent. Stoch. Process. their Appl. 13, 235–248. doi:10.1016/0304-4149(82)90011-4.

Klinkenberg, D., Backer, J.A., Didelot, X., Colijn, C., Wallinga, J., 2017. Simultaneous inference of phylogenetic and transmission trees in infectious disease outbreaks. PLoS Comput. Biol. 13, e1005495. doi:10.1371/journal.pcbi.1005495.

Krone, S.M., Neuhauser, C., 1997. Ancestral processes with selection. Theor. Popul. Biol. 51, 210–237. doi:10.1006/tpbi.1997.1299.

Li, Q., Scornavacca, C., Galtier, N., Chan, Y.B., 2021. The multilocus multispecies coalescent: a flexible new model of gene family evolution. Syst. Biol. 70, 822–837. doi:10.1093/sysbio/syaa084.

Maddison, W.P., 1997. Gene trees in species trees. Syst. Biol. 46, 523–536. doi:10.1093/sysbio/46.3.523.

Maddison, W.P., Knowles, L.L., 2006. Inferring phylogeny despite incomplete lineage sorting. Syst. Biol. 55, 21–30. doi:10.1080/10635150500354928.

Mallo, D., de Oliveira Martins, L., Posada, D., 2016. SimPhy: phylogenomic simulation of gene, locus and species trees. Syst. Biol. 65, 334–344. doi:10.1093/sysbio/syv082.

McVean, G.A.T., Cardin, N.J., 2005. Approximating the coalescent with recombination. Phil. Trans. R. Soc. B 360, 1387–1393. doi:10.1098/rstb.2005.1673.

Moran, P.A.P., 1958. Random processes in genetics. Math. Proc. Cambridge Philos. Soc. 54, 60–71. doi:10.1017/S0305004100033193.

Nee, S., Holmes, E.C., Rambaut, A., Harvey, P.H., 1995. Inferring population history from molecular phylogenies. Philos. Trans. R. Soc. Lond. B. Biol. Sci. 349, 25–31. doi:10.1098/rstb.1995.0087.

Nordborg, M., 1998. On the probability of Neanderthal ancestry. Am. J. Hum. Genet. 63, 1237–1240. doi:10.1086/302052.

Notohara, M., 1990. The coalescent and the genealogical process in geographically structured population. J. Math. Biol. 29, 59–75. doi:10.1007/BF00173909.

Palacios, J.A., Minin, V.N., 2012. Integrated nested Laplace approximation for Bayesian nonparametric phylodynamics, in: Uncertain. Artif. Intell. - Proc. 28th Conf. UAI 2012, pp. 726–735.

Paradis, E., Schliep, K., 2019. Ape 5.0: An environment for modern phylogenetics and evolutionary analyses in R. Bioinformatics 35, 526–528. doi:10.1093/bioinformatics/bty633.

Pybus, O.G., Rambaut, A., Harvey, P.H., 2000. An integrated framework for the inference of viral population history from reconstructed genealogies. Genetics 155, 1429–1437. doi:10.1093/genetics/155.3.1429.

Rabiner, L., 1989. A tutorial on hidden Markov models and selected applications in speech recognition. Proc. IEEE 77, 257–286. doi:10.1109/5.18626.

Rambaut, A., 2000. Estimating the rate of molecular evolution: incorporating non-contemporaneous sequences into maximum likelihood phylogenies. Bioinformatics 16, 395–399. doi:10.1093/bioinformatics/16.4.395.

Rasmussen, M.D., Hubisz, M.J., Gronau, I., Siepel, A., 2014. Genome-wide inference of ancestral recombination graphs. PLoS Genet. 10. doi:10.1371/journal.pgen.1004342.

Rasmussen, M.D., Kellis, M., 2012. Unified modeling of gene duplication, loss, and coalescence using a locus tree. Genome Res. 22, 755–765. doi:10.1101/gr.123901.111.

Rosenberg, N.A., Nordborg, M., 2002. Genealogical trees, coalescent theory and the analysis of genetic polymorphisms. Nat. Rev. Genet. 3, 380–90. doi:10.1038/nrg795.

Slatkin, M., Hudson, R.R., 1991. Pairwise comparisons of mitochondrial DNA sequences in stable and exponentially growing populations. Genetics 129, 555–562. doi:10.1093/genetics/129.2.555.

Takahata, N., Nei, M., 1985. Gene genealogy and variance of interpopulational nucleotide differences. Genetics 110, 325–344. doi:10.1093/genetics/110.2.325.

Tavaré, S., 1984. Line-of-descent and genealogical processes, and their applications in population genetics models. Theor. Popul. Biol. 26, 119–164. doi:10.1016/0040-5809(84)90027-3.

Volz, E.M., 2012. Complex population dynamics and the coalescent under neutrality. Genetics 190, 187–201. doi:10.1534/genetics.111.134627.

Volz, E.M., Didelot, X., 2018. Modeling the growth and decline of pathogen effective population size provides insight into epidemic dynamics and drivers of antimicrobial resistance. Syst. Biol. 67, 719–728. doi:10.1093/sysbio/syy007.

Volz, E.M., Kosakovsky Pond, S.L., Ward, M.J., Leigh Brown, A.J., Frost, S.D.W., 2009. Phylodynamics of infectious disease epidemics. Genetics 183, 1421–1430. doi:10.1534/genetics.109.106021.

Wakeley, J., 2009. Coalescent theory: an introduction. Roberts and Company Publishers.

Wright, S., 1931. Evolution in Mendelian populations. Genetics 16, 97–159. doi:10.1093/genetics/16.2.97.

Wymant, C., Hall, M., Ratmann, O., Bonsall, D., Golubchik, T., de Cesare, M., Gall, A., Cornelissen, M., Fraser, C., 2018. PHYLOSCANNER: Inferring transmission from within-and between-host pathogen genetic diversity. Mol. Biol. Evol. 35, 719–733. doi:10.1093/molbev/msx304.

Zucchini, W., MacDonald, I.L., 2009. Hidden Markov models for time series: an introduction using R. Chapman and Hall/CRC.

